# Metabolomic Network Analysis Reveals Reorganization of Lipid and Steroid Programs Linked to Right Ventricular-Pulmonary Vascular Function in Pulmonary Hypertension

**DOI:** 10.64898/2026.06.16.732773

**Authors:** Ivor W. Clinton, Julie C. Coursen, Darin T. Rosen, Karthik Suresh, Aparna Balasubramanian, Todd M. Kolb, Rachel L. Damico, Stephen C. Mathai, Steven Hsu, Monica Mukherjee, J. Emanuel Finet, Gabriele Grunig, John Barnard, Anna R. Hemnes, Jane A. Leopold, Evelyn M. Horn, Erika B. Rosenzweig, Franz Rischard, Robert P. Frantz, Serpil C. Erzurum, Wendy King, Gerald J. Beck, Nicholas S. Hill, Paul M. Hassoun, Catherine E. Simpson, PVDOMICS Study Group

**Author notes:** **Corresponding Author:** Catherine E. Simpson, MD, MHS, 1830 East Monument Street, 5th Floor, Baltimore, MD, USA 21205.

## Abstract

**Background:** Pulmonary arterial hypertension (PAH) is characterized by circulating metabolic alterations, but whether these reflect disease-specific metabolic programs or reorganization of normal metabolic architecture, and how they relate to right ventricular-pulmonary vascular function (RV-PV), remains unclear. We hypothesized that the PAH metabolome is organized into biologically coherent, co-regulated metabolic modules whose relationships to RV-PV function would provide insight into known and novel metabolic pathways.

**Methods:** We applied weighted gene co-expression network analysis (WGCNA) to untargeted metabolomic data from 412 PAH patients enrolled in the multicenter PVDOMICS study. Module preservation analysis was performed in 85 healthy controls, with external replication in an independent single-center pulmonary hypertension cohort of 89 patients.

**Results:** WGCNA identified 16 distinct metabolic modules organized around biologically coherent programs. A coherent fatty acid axis, spanning substrate pools, β-oxidation intermediates, and conjugated fatty acid disposal products, formed a central organizing structure, with downstream fatty acid oxidation modules strongly associated with adverse hemodynamics and worse RV-pulmonary artery (PA) coupling. Acylcholine-enriched and 5α-reduced androgen metabolite modules were associated with favorable hemodynamic indices. Module architecture was largely preserved in healthy controls, with subtle disease-associated modular reorganization, rather than emergence of novel modules, observed in PAH. Core modules were recovered in the replication cohort with conserved hub metabolites.

**Conclusions:** These findings establish a systems-level framework demonstrating that PAH involves structured intensification and reorganization of interconnected metabolic programs associated with favorable and adverse RV-PV phenotypes. This work provides new insight into the metabolic architecture underlying PAH and identifies coordinated metabolic pathways linked to pulmonary vascular and right ventricular function.

## INTRODUCTION

Pulmonary arterial hypertension (PAH) is a progressive disease hallmarked by elevated pulmonary artery (PA) pressures and pulmonary vascular remodeling resulting in right ventricular (RV) failure if untreated.^1–3^ Patient outcomes are determined primarily by the RV’s ability to adapt to increasing afterload.^1–3^ Altered metabolism is a well-established feature of PAH;^1–3^ however, organization of the PAH metabolome and its relationship to RV-pulmonary vascular (PV) physiology remain incompletely understood. While numerous studies have identified alterations in circulating metabolites in PAH, including perturbations in fatty acid metabolism, amino acid handling, and steroid biology, these observations have largely been interpreted at the level of individual metabolites or pathways.^4–8^ Whether metabolic alterations observed in PAH represent disease-specific programs or a reorganization of normal metabolic architecture has not been systematically addressed.

In this study, we applied weighted gene co-expression network analysis (WGCNA) to untargeted metabolomic data to define biologically coherent modules of co-expressed metabolites.^9–12^ By first identifying network-level metabolic programs and then interrogating their relationships to RV-PV physiology, we aimed to move beyond single-metabolite associations toward a systems-level understanding of metabolic organization in PAH. Through integration of multicompartment sampling across the RV-PV circuit, module preservation analysis in healthy controls (HC), and replication of results in an external cohort, we sought to distinguish stable metabolic architecture from disease-associated reorganization, and to identify coordinated programs that may underlie adaptive versus maladaptive responses to chronic pressure overload. We developed an interactive atlas to facilitate exploration of WGCNA-derived metabolite modules, hub metabolites, RV-PV associations, and cross-cohort replication.

## METHODS

### Discovery Cohort (PVDOMICS)

Our discovery cohort included 412 patients with PAH from the multicenter Pulmonary Vascular Disease Phenomics (PVDOMICS) study. The PVDOMICS study cohort included subjects with all forms of PH who underwent comprehensive hemodynamic assessment, RV imaging with echocardiography and cardiac MRI (CMR), and metabolomic profiling. Comparator patients with PH-predisposing conditions, but without PH, and HC were also enrolled. Procedures for enrollment, methods, and clinical testing have been previously published.^13,14^ To limit potential metabolic heterogeneity related to different forms of pulmonary vascular disease, we analyzed the metabolomes of PVDOMICS subjects cleanly adjudicated as having PAH.

### Replication Cohort (CALIPSO)

To evaluate the replicability of metabolic modules discovered in PVDOMICS, we utilized our single-center CALIPSO cohort. The CALIPSO cohort included 89 patients from a single tertiary-care center (Johns Hopkins Hospital) who were enrolled between 2013 and 2022 after referral for suspected PH. No patients in CALIPSO were co-enrolled in PVDOMICS. The CALIPSO protocol was approved by the Johns Hopkins Institutional Review Board and has been previously published, and all patients provided informed consent.^15–18^

### Metabolic Profiling

Circulating plasma samples from both discovery and replication cohorts were obtained during RHC under fasting conditions. Samples were stored at -80 °C and remained frozen until processing. Mass-spectrometry based metabolomic profiling, quality control, and metabolite identification were performed by Metabolon, Inc. using their Global HD4 platform for all PVDOMICS and CALIPSO samples. Further methodological details are provided in Supplemental Methods.

### WGCNA Construction

Weighted gene co-expression network analysis (WGCNA) was performed using the *wgcna* package in R to identify modules of co-expressed metabolites. Modules were labeled using standard WGCNA color assignments. Modules were biologically annotated by integrating Metabolon-provided metabolite subclass designations with predominant metabolite composition, known biochemical pathways, and hub metabolites, with annotations assigned by consensus review. Further details regarding WGCNA network construction are provided in Supplemental Methods.

### Module-Trait Associations

Module eigenmetabolites were calculated to summarize module-level variation. Module eigenmetabolites represent dimensionless latent variables derived from principal component analysis of normalized metabolite abundances. Module coherence was assessed by the proportion of variance explained by the eigenmetabolite (MEVar). Intramodular connectivity was quantified using kME, defined as the correlation between each metabolite and its module eigenmetabolite, and was used to identify hub metabolites.

In both discovery and replication cohorts, module eigenmetabolites were evaluated for association with invasive hemodynamic measurements obtained during right heart catheterization (RHC) and RV imaging using multivariable linear regression adjusted for age, sex, and BMI. Effect estimates from regression models were scaled to reflect differences per interquartile range of the module eigenmetabolite and are presented as β/SE [95% confidence interval]. Module gradients and projections are presented on the native eigenmetabolite scale. Additional models incorporated interaction terms to evaluate whether associations between module eigenmetabolites and hemodynamic traits differed between incident and prevalent disease states.

### Projection into Healthy Controls

To examine whether PAH-derived modules are disease-specific versus representative of normal metabolic architecture, we projected each module’s eigenmetabolite into the metabolomes of 85 HC enrolled in PVDOMICS using loadings derived from PAH patients. Module preservation in HC was evaluated by comparing eigenmetabolite means and medians to assess global module shift, measuring kME (metabolite-eigenmetabolite correlation) to assess module hub preservation, and examining principal component loadings correlations to assess preservation of module structure.

### Gradient Analysis

In PVDOMICS, availability of blood samples collected from multiple sites in the pulmonary circulation and from peripheral veins enabled additional analysis of transpulmonary eigenmetabolite gradients as well as comparisons of pulmonary versus peripheral venous eigenmetabolite abundance. Eigenmetabolite abundances per vascular compartment (mixed venous and wedge) and peripheral venous compartment were quantified and compared for each module. Full details of transpulmonary gradient analysis are provided in Supplemental Methods.

## RESULTS

### Metabolic Architecture in the PVDOMICS Cohort

Baseline demographic and hemodynamic characteristics for the 412 patients studied from the PVDOMICS cohort are summarized in Table 1. Average age (mean ± SD) was 54 ± 14.5 years old, 71.8% were women, and 73.8% were White. Baseline right atrial (RA) pressure was 7.2 ± 4.9 mmHg, mean PA pressure (mPAP) 42.6 ± 14.6 mmHg, pulmonary vascular resistance (PVR) 6.8 ± 4.5 Wood units (WU), and RVEF 39 ± 12%. RV-PA coupling surrogate TAPSE/PASP was 0.34 ± 0.18 mm/mmHg.

**Table 1.** Baseline Demographic and Hemodynamic Characteristics

| **Variable** | **Discovery Cohort**<br>**(n=412)** | **Validation Cohort**<br>**(n=89)** | **p-value** |
| --- | --- | --- | --- |
| Age, years | 54.0 (14.5) | 57.1 (13.3) | 0.06 |
| BMI, kg/m<sup>2</sup> | 29.1 (7.6) | 29.1 (6.8) | 0.98 |
| Sex, n (%female) | 296 (71.8%) | 72 (82.8%) |  |
| Race, n (%white) | 304 (73.8%) | 62 (74.7%) |  |
| RA mean, mmHg | 7.2 (4.9) | 6.4 (3.9) | 0.07 |
| mPAP, mmHg | 42.6 (14.6) | 31.4 (13.1) | <0.01 |
| PCWP, mmHg | 11.1 (5.6) | 10.3 (4.3) | 0.14 |
| PVR, Woods units | 6.8 (4.5) | 5.3 (4.3) | <0.01 |
| Cardiac Output,<br>L/min | 5.0 (1.7) | 4.6 (1.3) | 0.06 |
| Cardiac Index,<br>L/min/m<sup>2</sup> | 2.6 (0.8) | 2.5 (0.7) | 0.16 |
| RVEF, % | 39.0 (12.0) | — | — |
| TAPSE, mm | 18.8 (4.5) | — | — |
| TAPSE/PASP,<br>mm/mmHg | 0.34 (0.18) | — | — |
| Ees/Ea | — | 1.11 (0.62) | — |

Data are means (SD) or n (%) as appropriate. TAPSE/PASP values above 0.31 mm/mmHg are indicative of preserved coupling.<sup>39</sup> BMI: body mass index; RA: right atrial; mPAP: mean pulmonary artery pressure; PCWP: pulmonary capillary wedge pressure; PVR: pulmonary vascular resistance; TAPSE: tricuspid annular plane systolic excursion; PASP: pulmonary artery systolic pressure; Ees: end systolic elastance; Ea: arterial elastance.

WGCNA generated 16 distinct modules with low inter-module correlation, supporting inter-module distinctiveness, and high within-module variance explained by PC1, supporting intramodular coherence for the majority of modules (Figure 1, Supplemental Table 1). Examination of eigenmetabolite dendrogram structure together with module metabolite assignments and hubs revealed several biologically coherent metabolic programs summarized in Table 2. A fatty acid (FA) axis spanning three different modules was observed (brown, blue and green modules). Steroid metabolism was split into two related modules (yellow and cyan), and bile acid metabolism was captured across multiple modules, including primary conjugated (salmon) and secondary bile acids (tan). Full metabolite-module assignments, with hubs delineated by kME, are provided in Supplemental Table 2.

**Figure 1.**
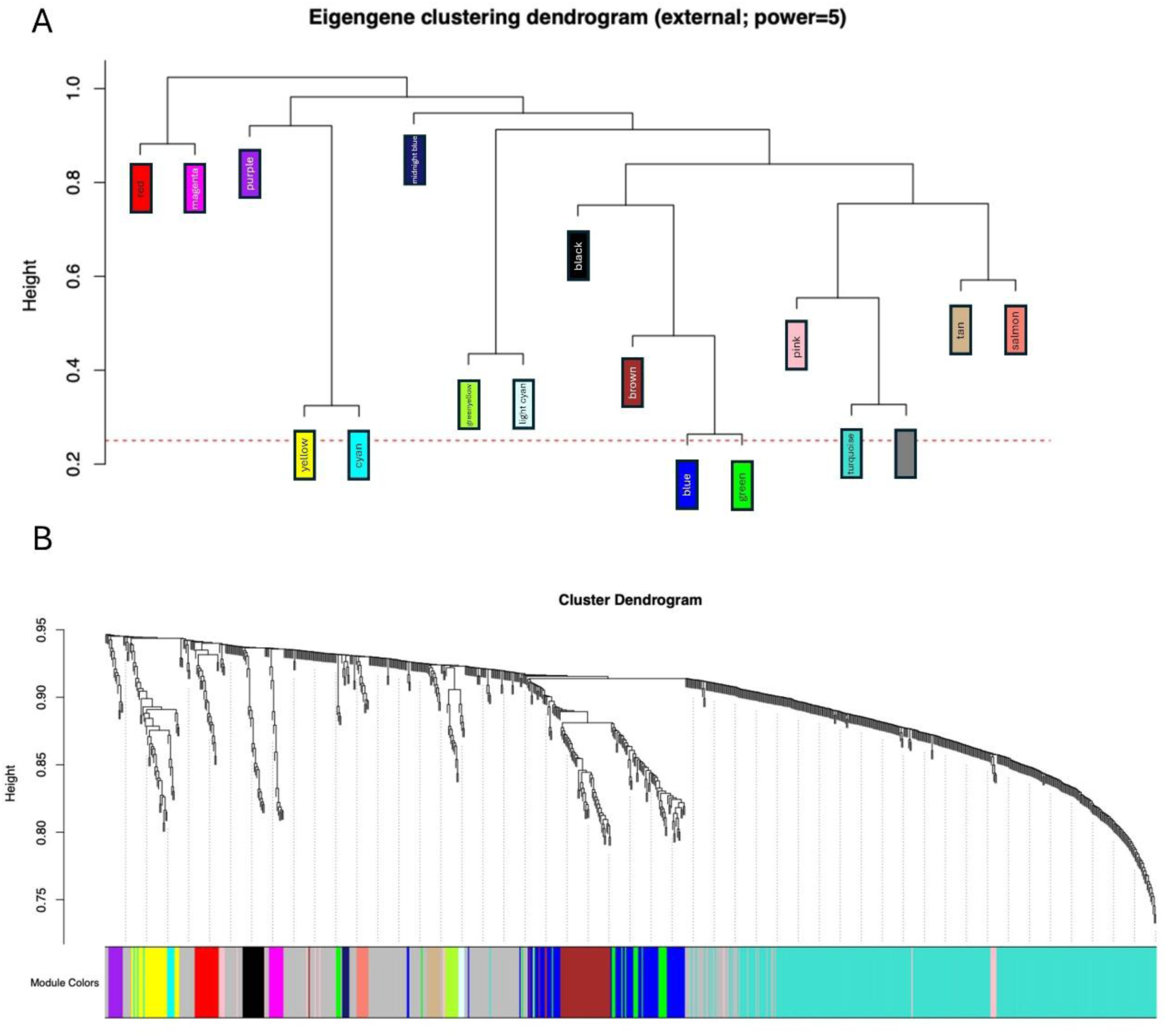
Modular metabolic architecture of the PAH metabolome defined by weighted gene co-expression network analysis (WGCNA). A: Eigenmetabolite clustering dendrogram illustrating hierarchical relationships among 16 co-expression modules identified by WGCNA from the PVDOMICS discovery cohort. Each colored block represents a module eigenmetabolite (first principal component). Branch height (y-axis) reflects inter-module similarity, with modules merging at lower heights sharing greater metabolomic similarity. Closely related modules include the fatty acid (FA) oxidation axis (blue, green, and brown), the adrenal steroid axis (yellow and cyan), and the primary and secondary bile acid programs (salmon and tan). B: Cluster dendrogram of all 974 metabolites retained after quality control filtering. Each branch represents an individual metabolite, and the color bar beneath the dendrogram indicates module assignment. Module colors correspond to standard WGCNA color labels. Grey denotes unassigned metabolites.

**Table 2.** Discovery Cohort Summary of Metabolic Programs

| ME | Color | Metabolite Summary |
| --- | --- | --- |
| ME1 | turquoise | Modified nucleosides, N-acetylated amino acids, short- and medium-chain acylcarnitines, kynurenine pathway metabolites, methionine/cysteine metabolites |
| ME2 | blue | Medium and long-chain acylcarnitines, dicarboxylic acids, hydroxy fatty acids, ketone bodies, and TCA cycle intermediates |
| ME3 | brown | Odd-, long-, medium-chain fatty acids (saturated, monounsaturated, and polyunsaturated) |
| ME4 | yellow | Sulfated and glucuronidated pregnenolone, androstenediol steroids, DHEA-S, and DHEA-S derivatives |
| ME5 | green | Acyl-glycine and acyl-glutamine conjugates with short-chain hydroxyacylcarnitines |
| ME6 | red | Phospholipid head groups, sphingoid bases, sphingosine-1-phosphate, glycerophosphoserines, and polyamines |
| ME7 | black | Bilirubin, biliverdin, and multiple bilirubin degradation products |
| ME8 | pink | Carboxyethylated amino acids, pyruvate, lactate, and branched-chain α-hydroxy acids |
| ME9 | magenta | Long-chain fatty acylcholines (palmitoyl-, oleoyl-, stearoyl-, arachidonoylcholine) with glycerophosphocholine |
| ME10 | purple | Primarily composed of unidentified metabolites. Known metabolites include dicarboxylic acids (C20-DC, C20:1-DC) and hydroxylaurate |
| ME11 | greenyellow | Xanthine metabolism: caffeine, paraxanthine, and downstream methylurates |
| ME12 | tan | Secondary bile acid conjugates: glycine-, taurine-, and sulfate-conjugated lithocholate and deoxycholate derivatives |
| ME13 | salmon | Glycine- and taurine-conjugated primary bile acids |
| ME14 | cyan | 5α-Reduced androgen metabolites (epiandrosterone sulfate, androsterone sulfate, 5α-androstanediol sulfates) |
| ME15 | midnightblue | Ursodeoxycholate, isoursodeoxycholate, and their glycine/sulfate conjugates |
| ME16 | lightcyan | Xanthine metabolism: theobromine, 7-methylxanthine, 3-methylxanthine, and downstream methylurates |

### 1) Fatty Acid Substrate Pools and Products of FA Oxidation

Blue, brown, and green modules emerged as closely related FA metabolism programs. The blue module was primarily enriched in long-chain acylcarnitines and dicarboxylic (DC) fatty acids, products of mitochondrial β-oxidation and peroxisomal ω-oxidation, respectively.^19^ The green module contained acyl-glycine and acyl-glutamine conjugates, and hydroxyacylcarnitines, intermediate and downstream disposal products of fatty acid β-oxidation.^20^ The brown module was enriched with odd-chain fatty acids (OCFA), long chain unsaturated fatty acids, and several medium chain fatty acids, all substrates for β-oxidation.^21^

Both blue and green FA oxidation modules were strongly associated with adverse hemodynamics. Per IQR increase in the blue module eigenmetabolite, PVR increased by 1.69 WU [1.12–2.26] (q = 8.25 × 10^-7^) and mPAP increased by 6.24 mmHg [4.42–8.06] (q = 1.18 × 10^-8^). Per IQR increase in the green module eigenmetabolite, PVR increased by 1.26 WU [0.69–1.83] (q = 3.79 × 10^-4^) and mPAP increased by 4.51 mmHg [2.66–6.36] (q = 6.60× 10^-5^). Comprehensive module-trait association results are provided in Figures 2 and 3. Hemodynamic trait associations for blue and green modules persisted with fluid challenge and vasodilator challenge (Supplemental Table 3). Furthermore, both blue and green modules were associated with lower RVEF by CMR (blue: -2.05%/IQR [-4.40– -0.76], q = 3.3 × 10^-2^; green: -2.61%/IQR [-4.40– -0.83], q =2.7 × 10^-2^). In contrast, the brown FA substrate pool module was not associated with elevated right sided pressures globally, but did demonstrate narrower associations with mPAP (2.47 mmHg/IQR [0.66–4.27], q = 3.7× 10^-2^) and PVR (0.96 WU/IQR [0.40–1.52], q = 7.9 × 10^-3^) that persisted during vasodilator oxygen challenge (Supplemental Table 3).

**Figure 2.**
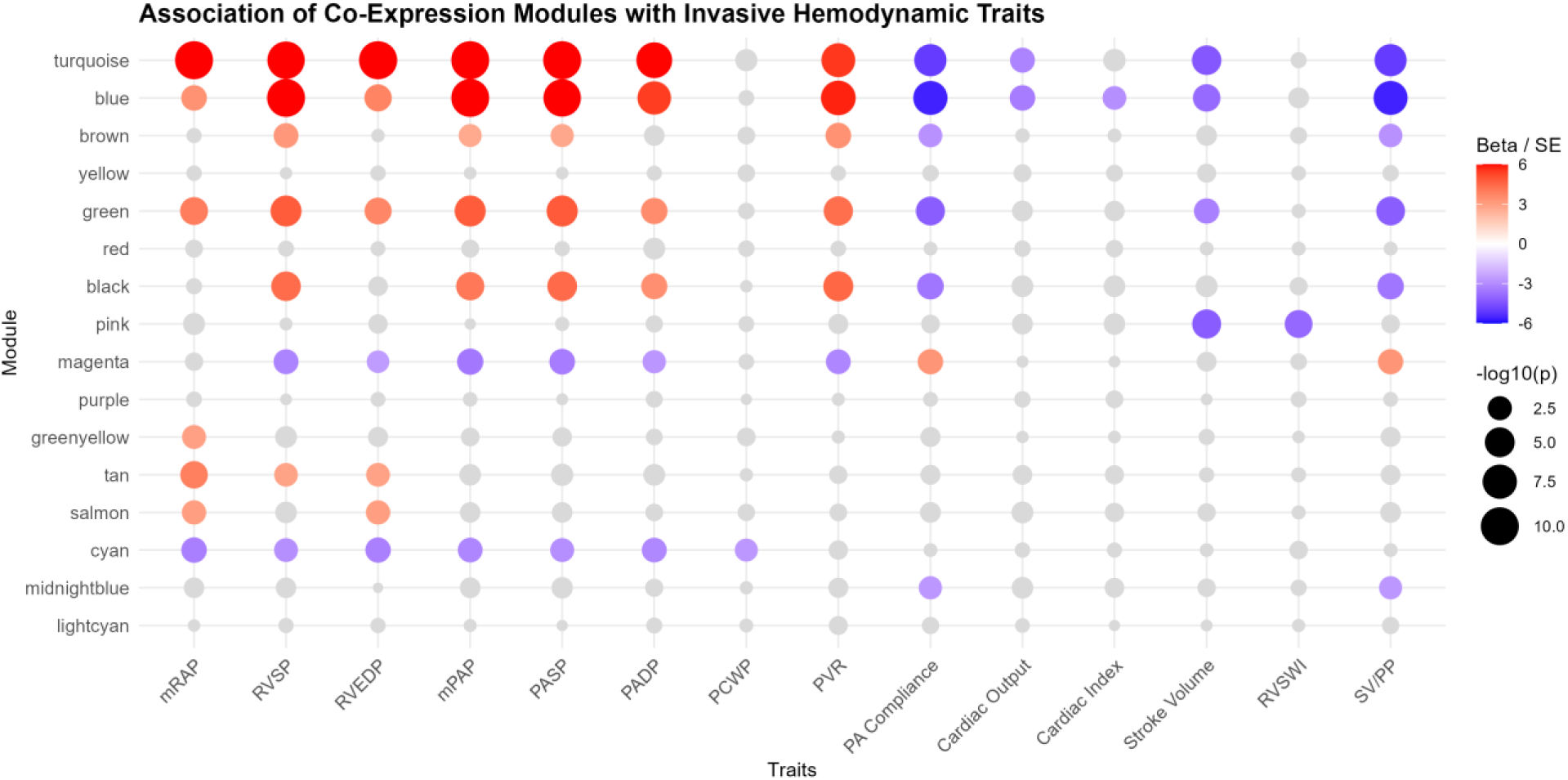
Module-trait association matrix for metabolite modules and invasive hemodynamic traits from the discovery cohort. Bubble plot displaying associations between module eigenmetabolites (y-axis) and hemodynamic traits (x-axis) from multivariable linear regression models adjusted for age, sex, and BMI. Bubble color indicates the direction and magnitude of the association (β/SE), with red indicating positive and blue indicating negative associations. Bubble size reflects statistical significance, scaled by −log₁₀(*p*). Traits include mRAP (mean right atrial pressure, mmHg), RVSP/RVEDP (RV systolic and end-diastolic pressures, mmHg), PASP/PADP (pulmonary artery systolic and diastolic pressures, mmHg), mPAP (mean pulmonary artery pressure, mmHg), PCWP (pulmonary capillary wedge pressure, mmHg), cardiac output (L/min), cardiac index (L/min/m^2^), stroke volume (mL), PVR (pulmonary vascular resistance, Wood units), RVSWI (RV stroke work index, g·m/m²), SV/PP (stroke volume to pulse pressure ratio, mL/mmHg).

**Figure 3.**
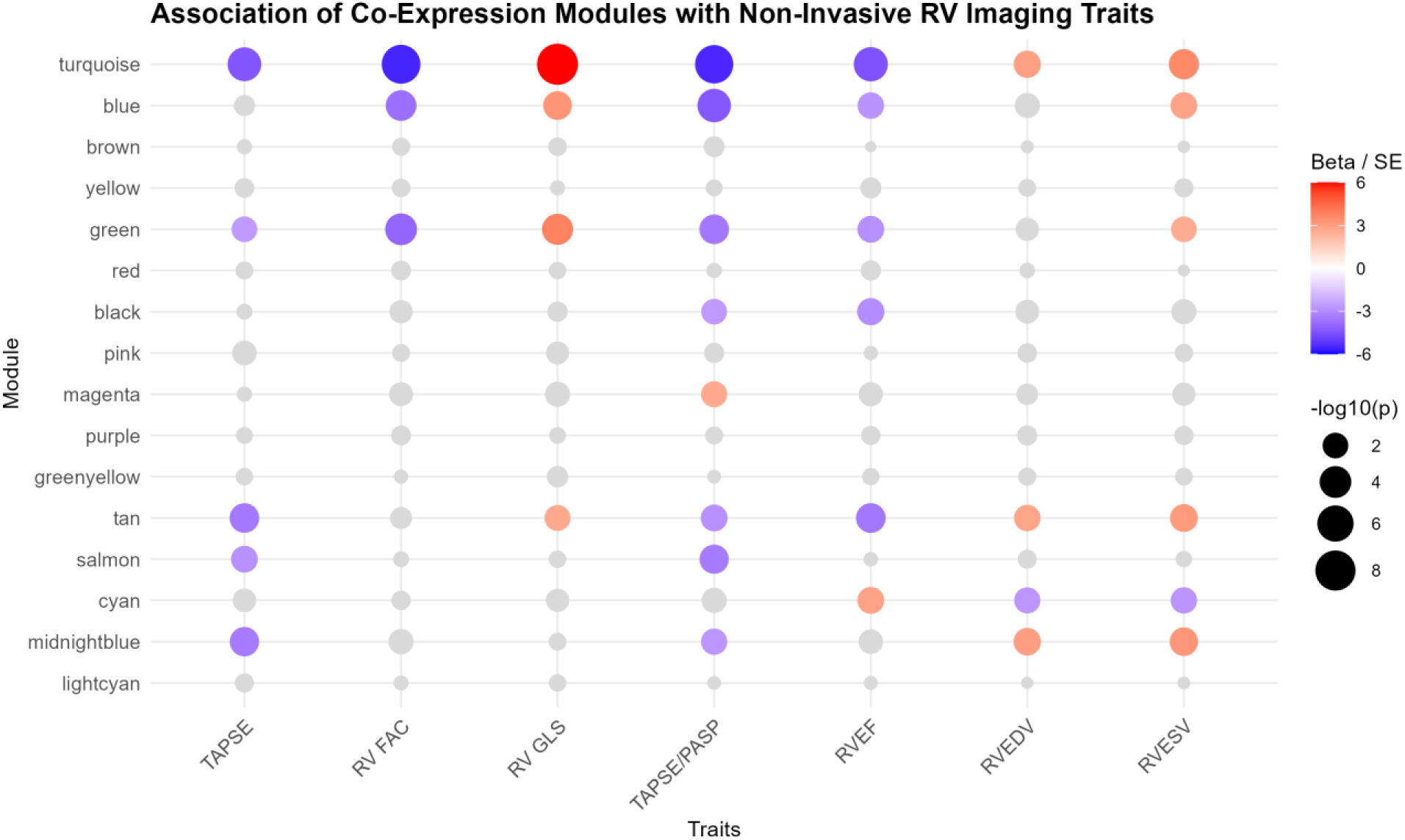
Module-trait association matrix for metabolite modules (y-axis) and echocardiographic and CMR-derived traits (x-axis) from the discovery cohort. Associations were derived from multivariable linear regression models adjusted for age, sex, and BMI. Bubble color represents the direction and magnitude of association (β/SE), with red indicating positive and blue indicating negative associations. Bubble size corresponds to statistical significance, scaled by −log₁₀(p-value). Traits include TAPSE (tricuspid annular plane systolic excursion, mm), RV FAC (RV fractional area change, %), RV GLS (global longitudinal strain, %), TAPSE/PASP (ratio of TAPSE to pulmonary artery systolic pressure, mm/mmHg), RVEF (ejection fraction, %), RVEDV (RV end-diastolic volume, mL), RVESV (RV end-systolic volume, mL).

Blue and green FA oxidation modules, despite their associations with adverse hemodynamics, did not demonstrate a consistent gradient across the pulmonary circulation (Table 3).

**Table 3.**
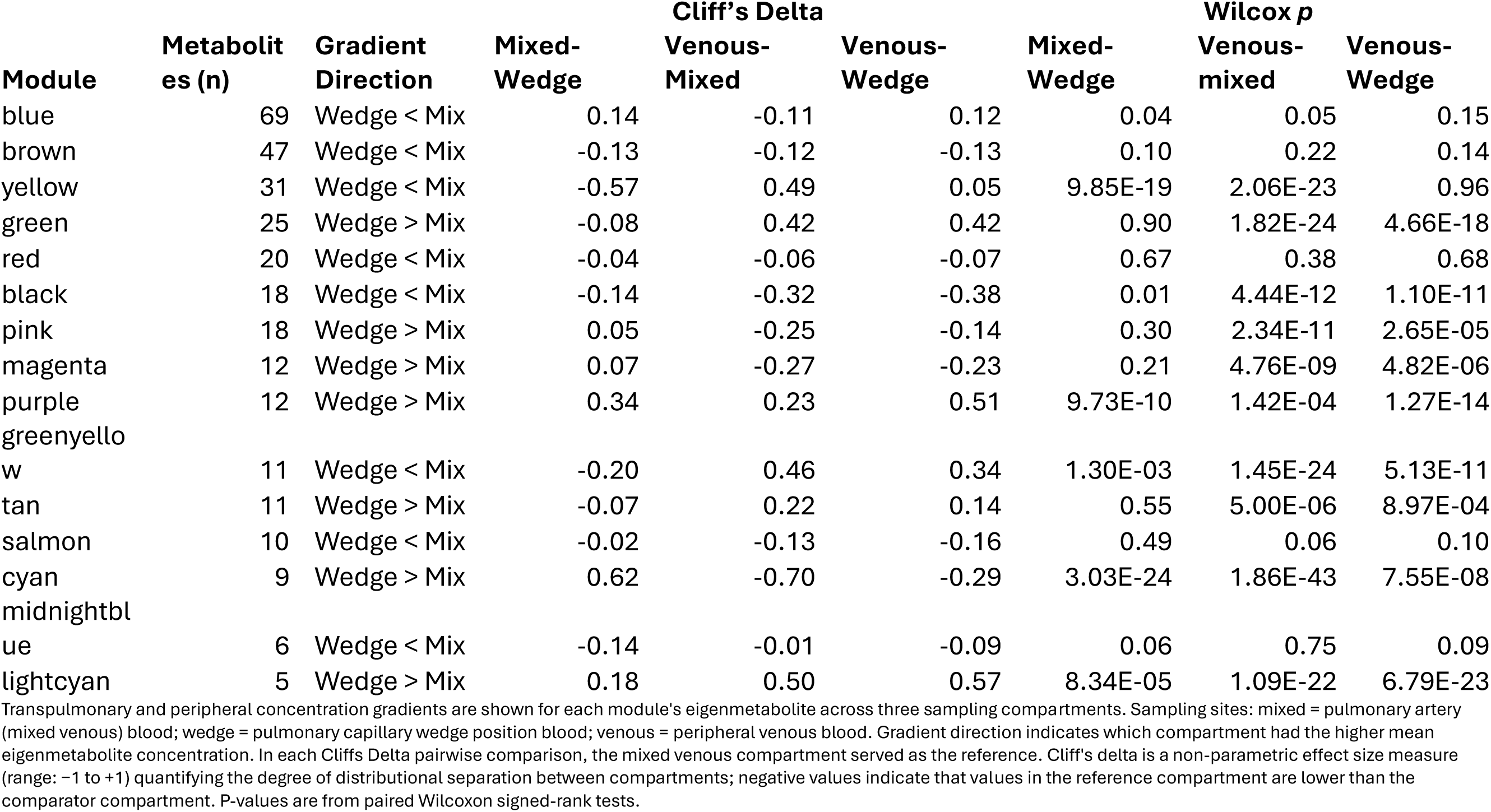
Transpulmonary Gradient Analysis in Discovery Cohort

### 2) Membrane Lipids and Acylcholines

A second lipid-enriched program was identified in modules red and magenta, which cluster together but branch distinctly in the dendrogram, clearly separated in space from the FA oxidation axis (Figure 1). The red module contained sphingoid bases, phosphorylated derivatives (sphingosine-1-phosphate), and glycerophosphoserines (GPS), constituents of sphingolipid metabolism and membrane phospholipid signaling.^22^ Magenta was highly internally coherent (MEVar 0.75) and composed almost exclusively of long-chain acylcholines (palmitoylcholine, oleocholine, stearoylcholine) and glycerophosphorylcholine (GPC). In contrast to FA oxidation modules, this magenta acylcholine module was associated with lower mPAP (-3.65 mmHg/IQR [-5.69– -1.62], q = 5.1 × 10^-3^), with stability of that association with fluid challenge and vasodilator challenge (Supplemental Table 3). In gradient analysis, the acylcholine module demonstrated significant eigenmetabolite reduction in the periphery, with peripheral venous abundances significantly lower than mixed venous (Cliff’s delta = -0.27, *p* = 4.76 × 10^-9^) or wedged abundances (Cliff’s delta = -0.23, *p* = 4.82 × 10^-6^), without change in eigenmetabolite abundance from mixed venous to wedged positions in the overall PAH cohort. In prevalent-only analyses, however, there was a significant transpulmonary release pattern, with eigenmetabolite mean gradient 0.42, *p* = 0.004. There was a trend toward a transpulmonary uptake pattern in incident PAH, with eigenmetabolite mean gradient of -0.79, the largest mean gradient observed among all module-specific compartment comparisons made in our analyses; however, there were few incident PAH patients with multicompartment sampling (n = 17), and the *p* value is not significant at 0.14. There was a difference in incident vs prevalent transpulmonary gradients for the magenta module by formal interaction testing, with *p* = 0.039.

In contrast to the acylcholine module, the red module comprised membrane signaling lipids was not gradient-active, and the eigenmetabolite was not significantly associated with hemodynamics or RV functional traits.

### 3) Steroid Metabolism

Both yellow and cyan modules were enriched in adrenal-derived steroids. Yellow was composed of sulfated sex hormone precursors, including DHEA-S, pregnenolone sulfate, and related C21 intermediates.^23^ In contrast, cyan was enriched with sulfated, 5α-reduced, steroid derivatives of DHEA, including androsterone sulfate, and androstenediol sulfates/disulfates.^24^ Similarity between these modules was reflected by their low branch height, yet cyan is distinguished by its high internal coherence, with MEVar 0.71. While the steroid program that contains DHEA-S (yellow) had no significant associations with RHC or CMR derived indices, the sulfated testosterone derivative program (cyan) was associated with lower mPAP (-3.80 mmHg/IQR [-6.17– -1.43], q = 1.3 × 10^-2^) and higher RVEF (3.51%/IQR [1.11 to 5.91], q = 2.7 × 10^-2^).

These two sex hormone modules were strikingly gradient-active, with highly significant shifts of substantial magnitude observed between mixed venous and wedged compartments, and, notably, in opposite directions (Table 3). The yellow module displayed a clear transpulmonary uptake pattern (Cliff’s delta = -0.57, *p* = 9.85 × 10^-19^) and this pattern remained directionally consistent in incident and prevalent cases. In contrast, the cyan/androgen metabolite module displayed a clear transpulmonary release pattern (Cliff’s delta = 0.62, *p* = 3.03 × 10^-24^), with peripheral tissues appearing to serve as a sink (that is, significantly lower abundances are observed in venous blood compared to wedged blood; Cliff’s delta = -0.70, *p* = 1.86 × 10^-43^). The transpulmonary pattern for cyan, similar to the yellow sex hormone module, remained directionally consistent in incident and prevalent cases.

### 4) Bile Acid Metabolism

Several modules captured distinct axes of bile acid biology. Salmon contained primary conjugated bile acids, tan contained secondary bile acids and sulfated conjugates, and midnightblue contained ursodeoxycholic acid and its derivatives. Salmon and tan branched together at mid-dendrogram height, indicating greater similarity to each other than to midnightblue. Consistent with this biologic similarity, both salmon and tan were associated with higher RA pressures (salmon: 0.97 mmHg/IQR [0.34–1.60], q = 1.9 × 10^-2^; tan: 1.24 mmHg/IQR [0.61–1.90], q = 1.8 × 10^-3^) and worse TAPSE/PASP (salmon: -0.05 mm/mmHg/IQR [-0.08– -0.02], q = 7.2 × 10^-3^ ; tan: -0.04 mm/mmHg/IQR [-0.07– -0.01], q = 2.8× 10^-2^). Only tan was associated with worse RVEF (-3.29%/IQR [-5.12 to -1.45], q = 5.4 × 10^-3^), while salmon had no CMR associations. None of the bile acid modules demonstrated eigenmetabolite shifts across the RV-PV circuit, however the trait-associated secondary bile acid module, tan, displayed significantly higher abundances in the periphery relative to mixed venous (Cliff’s delta = 0.22, *p* = 5.00 × 10^-6^) and wedged compartments (Cliff’s delta = 0.14, *p* = 8.97 × 10^-4^).

### 5) Turquoise Axis and Catabolic Byproducts

The turquoise module was a large module containing 344 metabolites with low internal coherence (MEVar 0.32). A complete list of turquoise metabolites is in Supplemental Table 1, but constituents of note include modified nucleosides, short- and medium-chain acylcarnitines, and kynurenine pathway metabolites. The turquoise module was robustly associated with adverse RV-PV measures, including higher RA mean (2.42 mmHg/IQR [1.76–3.09], q = 2.28 × 10^-9^), mPAP (6.96 mmHg/IQR [4.94–9.00], q = 1.18 × 10^-8^), PVR (1.81 WU/IQR [1.17–2.45], q = 2.71 × 10^-6^), worse TAPSE/PASP (-0.08 mm/mmHg/IQR [-0.11– -0.05], q = 3.74 × 10^-6^), and worse RVEF (-4.78%/IQR [-6.86– -2.70], q = 2.3 × 10^-4^). Adverse hemodynamics persisted with fluid challenge and vasodilator challenge (Supplemental Table 3).

### Module Preservation Analysis in Healthy Controls

Overall, metabolic architecture identified in PAH was largely preserved in HC with only subtle shifts observed across modules (Table 4). Most modules demonstrated preserved structure and hub concordance, with the brown FA substrate pool and adrenal steroid modules (yellow, cyan) showing particularly strong preservation. Notably, red and brown modules exhibited greater coherence when projected into HC than in the PAH cohort. Two modules deviated from this pattern, the green FA disposal module showed relative loss of structure and the turquoise module collapsed further in healthy controls (MEVar 0.32 → 0.15), such that it carries no summative, interpretable meaning in HC. The bile acid modules demonstrated preserved structure overall with variable hub concordance.

**Table 4.** Similarity Analysis of Discovery Cohort and Healthy Control Modules

| Module | MEVar<br>PAH | MEVar<br>HC | Loading<br>correlation | kME<br>correlation | TopK<br>Overlap | Wilcox<br>p-value | q-value |
| --- | --- | --- | --- | --- | --- | --- | --- |
| ME1 turquoise | 0.32 | 0.15 | 0.63 | 0.64 | 9 | 0.43 | 0.99 |
| ME2 blue | 0.42 | 0.36 | 0.87 | 0.89 | 12 | 0.78 | 0.99 |
| ME3 brown | 0.56 | 0.62 | 0.93 | 0.93 | 18 | 0.73 | 0.99 |
| ME4 yellow | 0.55 | 0.52 | 0.87 | 0.88 | 20 | 0.99 | 0.99 |
| ME5 green | 0.49 | 0.42 | 0.66 | 0.77 | 17 | 0.94 | 0.99 |
| ME6 red | 0.56 | 0.64 | 0.81 | 0.76 | 20 | 0.54 | 0.99 |
| ME7 black | 0.61 | 0.58 | 0.97 | 0.98 | 18 | 0.89 | 0.99 |
| ME8 pink | 0.39 | 0.42 | 0.95 | 0.96 | 18 | 0.93 | 0.99 |
| ME9 magenta | 0.75 | 0.71 | 0.94 | 0.96 | 12 | 0.93 | 0.99 |
| ME10 purple | 0.56 | 0.63 | 0.97 | 0.97 | 12 | 0.91 | 0.99 |
| ME11 greenyellow | 0.73 | 0.77 | 0.98 | 0.98 | 11 | 0.67 | 0.99 |
| ME12 tan | 0.58 | 0.42 | 0.92 | 0.89 | 11 | 0.83 | 0.99 |
| ME13 salmon | 0.64 | 0.51 | 0.98 | 0.97 | 10 | 0.88 | 0.99 |
| ME14 cyan | 0.71 | 0.61 | 0.94 | 0.98 | 9 | 0.87 | 0.99 |
| ME15<br>midnightblue | 0.69 | 0.67 | 0.93 | 0.62 | 6 | 0.89 | 0.99 |
| ME16 lightcyan | 0.86 | 0.89 | 0.93 | 0.84 | 5 | 0.99 | 0.99 |
| Modules were identified by WGCNA in the discovery cohort and projected onto healthy controls (HC). MEVar indicates the proportion of variance explained by each module's eigenmetabolite. Loading correlations reflect the Spearman correlation of metabolite loadings between cohorts. kME correlation reflects Spearman correlation of module membership values for hub metabolites between cohorts. TopK overlap indicates the number of shared metabolites among the top 20 hub metabolites between cohorts. Wilcoxon p-values compared eigenmetabolite scores between PAH and HC; q-values are FDR-adjusted. PAH: pulmonary arterial hypertension; HC healthy controls. |  |  |  |  |  |  |  |

### Reproducibility of Module Biology in Replication Cohort

To extend our replication efforts to an external cohort with more nuanced hemodynamic measurements, we analyzed the CALIPSO cohort. Average age (means ± SD) was 57.1 ± 13.3 years old, 82.8% were women, and 74.7% were White. Baseline RA mean 6.4 ± 3.9 mmHg, mPAP 31.4 ± 13.1 mmHg, and PVR 5.3 ± 4.5 WU. RV-PA coupling was derived from PV loops with an average Ees/Ea 1.11 ± 0.62.

WGCNA in the replication cohort identified 13 distinct metabolite modules (Supplemental Figures 1-2; Supplemental Table 4). Core metabolic programs identified in the discovery cohort were largely preserved (Table 5). The tripartite FA axis was recovered: substrate pool (brown → magenta, 63% overlap), acylcarnitine/DC acid (blue → yellow, 43% overlap), and conjugated disposal lipids (green → greenyellow, 64% overlap). Steroid modules mapped to a shared replication module (yellow and cyan → green, 77% and 100% overlap). Sphingolipid and acylcholine modules were moderately preserved (red → pink, 40% overlap; magenta → black, 58% overlap), and both embedded in more heterogenous modules encompassing broader lipid turnover programs. Bile acid modules were absorbed into the unassigned grey module (salmon/tan → grey, 82% and 90% overlap). Additionally, many of the same metabolites occupied hub positions in each module. For example, 10-heptadecenoate (discovery kME 0.89, replication kME 0.94) in the FA substrate pool, androstenediol (3α, 17β) monosulfate (0.92, 0.84) in the steroid axis, palmitoleoylcarnitine (0.89, 0.83) and dodecenedioate (0.74, 0.74) in the acylcarnitine/DC module, and palmitoylcholine in the acylcholine module (0.98,0.83). Complete replication cohort module-metabolite assignments with kMEs are provided in Supplemental Table 5. The catch-all turquoise module demonstrated high overlap of metabolites (68% overlap) with the discovery cohort turquoise model, again comprised of kynurenine pathway metabolites, modified nucleosides, and catabolic products.

**Table 5.** Similarity Analysis of Discovery Cohort and Analogous Replication Cohort Modules

| Discovery (A) | Replication (B) | Total Metabolites (A) | Total Metabolites (B) | Intersection | Union | Jaccard | Overlap |
| --- | --- | --- | --- | --- | --- | --- | --- |
| Turquoise | Turquoise | 344 | 406 | 236 | 514 | 0.46 | 0.69 |
| Blue | Yellow | 69 | 49 | 21 | 97 | 0.22 | 0.43 |
| Brown | Magenta | 47 | 27 | 17 | 57 | 0.30 | 0.63 |
| Yellow | Green | 31 | 40 | 24 | 47 | 0.51 | 0.77 |
| Green | Greenyellow | 25 | 11 | 7 | 29 | 0.24 | 0.64 |
| Red | Pink | 20 | 31 | 8 | 43 | 0.19 | 0.40 |
| Magenta | Black | 12 | 38 | 7 | 43 | 0.16 | 0.58 |
| Tan | Grey | 11 | 498 | 9 | 500 | 0.02 | 0.82 |
| Salmon | Grey | 10 | 498 | 9 | 499 | 0.02 | 0.90 |
| Cyan | Green | 9 | 40 | 9 | 40 | 0.23 | 1.00 |
| Midnightblue | Grey | 6 | 498 | 4 | 500 | 0.01 | 0.67 |
| WGCNA was performed independently in the discovery and replication cohorts. Each discovery module was matched to its best-corresponding replication module based on metabolite overlap. Discovery modules not shown were excluded due to small module size, predominance of unidentified metabolites, or lack of interpretable biological signal. Intersection, number of shared metabolites between modules; union, total number of unique metabolites across both modules; Jaccard index, ratio of intersection to union; overlap coefficient, ratio of intersection to the smaller module size. |  |  |  |  |  |  |  |

### Reproducibility of Module-Trait Analysis in Replication Cohort

We performed module-trait analysis on the OCFA and unsaturated FA substrate pool, adrenal steroid axis, and lipid signaling axes that were recapitulated in CALIPSO. In the full cohort, only one relatively weak association between the lipid substrate pool and lower RA pressure survived FDR correction (q = 0.02). Several nominal associations (*p* < 0.05) between the long-chain acylcholines module and lower pulmonary pressures were observed, directionally consistent with discovery cohort findings.

## DISCUSSION

Our findings demonstrate that RV-PV-associated metabolic programs represent disease-state intensification and alternative utilization of normal metabolic architecture, rather than emergence of novel, PAH-specific metabolic programs. Despite differences in cohort composition, sample size, and clinical context, we were able to re-demonstrate the core metabolic organization in an external replication cohort. These results indicate that the biologic organization captured by WGCNA was robust to differences in cohort structure, even as module boundaries shifted slightly with sample size and population heterogeneity.

A central component of the metabolic architecture identified in this study was a reproducible tripartite FA handling axis composed of a FA substrate pool, acylcarnitine/DC acids, and downstream conjugated disposal products. These modules formed a closely related structure in both cohorts, implying sequential handling of FA substrates across uptake, oxidation, and downstream disposal processes. Only the oxidation and conjugated disposal modules were associated with adverse hemodynamic and CMR indices, suggesting increased flux through FA pathways in disease, with incomplete oxidation and engagement of alternative disposal pathways. These findings align with prior observations of mitochondrial dysfunction and altered substrate utilization in the pressure-overloaded RV.^1–4^ Interestingly, despite strong hemodynamic associations observed with the FA oxidation modules, these modules were not clearly gradient-active, suggesting that shifts in FA handling may occur systemically in PAH, potentially via inter-organ cross-talk, rather than in processes isolated solely to the RV-PV unit.

The FA substrate pool module demonstrated less consistent associations with pulmonary pressures, though did demonstrate gradient sign reversal between incident and prevalent cases, with uptake in prevalent disease and a statistically significant interaction term. The substrate pool was comprised of several long-chain polyunsaturated FAs previously observed in PAH^4,25^, with linoleate and linolenate occupying top hub positions (kME 0.90 and 0.87, respectively), as well as omega-3 FAs EPA, DPA, and DHA, which are thought to exert anti-inflammatory and other protective effects in cardiovascular diseases.^26–28^ Other central hubs included monounsaturated and fully saturated odd chain FAs, such as 17-carbon heptadecanoate (kME 0.89), 19-carbon nonadecenoate (kME 0.84), and 15-carbon pentadecanoate (kME 0.82), which have not previously been associated with PAH. Transpulmonary uptake of these substrates in prevalent disease only may reflect compensatory fuel switching during disease progression. Notably, the terminal product of OCFA β-oxidation is propionyl-CoA, which can regenerate the TCA cycle under conditions when succinyl-CoA is depleted.^29,30^ We have previously shown TCA intermediates to be depleted in circulation and in RV tissue when the pressure-overloaded RV is dysfunctional in PH.^18^ Our findings suggest potential use of OCFA to resupply the TCA cycle as a means of adaptive substrate switching under stress.^1–3^ Direct assessment of substrate flux will be required to confirm the mechanistic hypotheses we propose here.

Other lipid-based modules, not related to oxidation, demonstrated favorable RV-PV associations. A module enriched for long-chain acylcholine species was associated with lower pulmonary pressures and improved indices of RV function. Although the biological role of acylcholines remains incompletely defined, their reproducible co-clustering with phospholipid intermediates suggests that they may reflect coordinated membrane-associated lipid processes. We have previously shown that higher serum concentrations were protective against development of PAH in patients with SSc, also suggesting a protective or adaptive role.^25^ Further studies are needed to clarify the role of acylcholines in PAH.

The other lipid-related module with favorable associations was the distal arm of the larger adrenal steroid axis. While prior work has demonstrated associations between DHEA-S and RV function, the module that contained DHEA-S was not associated with RV-PV measures in our analysis.^31,32^ In contrast, the downstream steroid module, which contained 5α-reduced androgen derivatives, demonstrated clear associations with lower right-sided pressures and improved RV-PA coupling. Gradient analysis showed both arms of the steroid axis to be highly gradient-active and contrasting, with uptake of precursor metabolites (yellow), and concurrent release of reduced androgen derivatives (cyan). This pattern is consistent with delivery and utilization of sex hormones across the RV-pulmonary circulation, in alignment with a robust body of preclinical work.^23,31,33,34^ Observed associations with circulating DHEA-S may reflect coordinated activity within this broader steroid metabolism program; our data showing preferential coupling of 5α-reduced androgen products to RV-PV indices (rather than sex hormone precursors) suggest that efficiency of key steroid conversion pathways may be more important to RV-PV function than the abundance of circulating prohormones like DHEA-S, and as such, conversion pathway constituents may prove to be more tractable therapeutic targets. Supporting this interpretation, the EDIPHY trial, a randomized, placebo-controlled crossover study of DHEA supplementation in PAH, showed no improvement in RV longitudinal strain and only modest, inconsistent effects across secondary endpoints despite robust increases in circulating DHEA-S.^35^

Bile acid (salmon and tan) and turquoise modules branched on the far right of the eigenmetabolite dendrogram, which became increasingly comprised of catabolic byproducts as the dendrogram progressed from left to right. These modules all demonstrated large increases in internal cohesion from HC to the PAH disease-state, suggesting that these programs undergo the most substantial disease-state reorganization. Hepatic congestion is one reasonable explanation for the observed associations between the primary and secondary bile acids modules and worse hemodynamic and RV functional measures. That said, there is emerging evidence suggesting bile acid metabolism may be intrinsic to the pulmonary circulation itself. Harvey et al. recently established a novel genetic and metabolic paradigm linking lysosomal dysfunction, NCOA7 deficiency, and transpulmonary oxysterol/bile acid metabolism to the pulmonary endothelium in PAH.^36^ However, in our analysis, we failed to detect primary bile acid transpulmonary gradients. Similarly, the strong association of the turquoise module with adverse hemodynamic and RV functional measures may reflect renal dysfunction due to congestion. Central hub metabolites from the turquoise module, pseudouridine, C-glycosyltryptophan, and hydroxyasparagine, are established correlates of measured glomerular filtration rate.^37^ Despite its heterogeneity representing many metabolic programs, increased turquoise coherence in PAH likely reflects alignment of disparate metabolic processes under systemic disease stress.

Taken together, our findings illuminate a reproducible, hierarchical structure to the human circulating metabolome that organizes into coordinated, differential associations with RV-PV physiology in PAH. By examining eigenmetabolites, weighted combinations of metabolites assigned to coherent modules, we are able to identify patterns of coordinated variation to show that PAH is associated with structured metabolic shifts in normal metabolic programs. Rather than identifying isolated perturbations in individual metabolites, we find shifts in interconnected metabolic systems, with sequential alterations in fatty acid handling, a potentially competing or protective membrane lipid and acylcholine signaling program and contrasting steroid metabolism axes. This systems-level organization provides a more interpretable framework for understanding metabolic remodeling in PAH than single-metabolite approaches and may better align with the complexity of RV-PV physiology.

Several limitations merit consideration. First, WGCNA identifies co-regulated metabolite modules based on correlation structure, which does not establish causal or mechanistic relationships.^38^ The biologic interpretations offered here are hypothesis-generating and require validation. Second, circulating metabolite abundances reflect the net balance of production, consumption, and clearance across multiple organ systems, and transpulmonary gradients cannot definitively localize metabolic activity to the pulmonary vasculature versus the RV. Third, the replication cohort was substantially smaller than the discovery cohort and enrolled patients with suspected PH. Though the majority were diagnosed with PAH, the cohort’s smaller size and broader composition limits statistical power for module-trait associations and introduces biologic heterogeneity. Last, though comprehensively profiled, the metabolome we analyze is incomplete, and relationships between metabolites may change as more metabolites are identified.

## Conclusion

In conclusion, this study defines the modular metabolic architecture of the circulating PAH metabolome and demonstrates that PAH is characterized by structured intensification of interconnected, co-regulated metabolic programs rather than the emergence of disease-specific pathways. Lipid metabolism, spanning FA handling, membrane lipid signaling, and steroid utilization, emerged as a central organizing axis. Multicompartment gradient analysis supports the pulmonary circulation as an active metabolic interface for steroid and, in a disease-stage-dependent manner, certain FA substrate pools. These findings provide a systems-level framework for understanding metabolic organization in PAH.

## Nonstandard Abbreviations

CMR: cardiac magnetic resonance imaging
DHEA-S: dehydroepiandrosterone sulfate
FA: fatty acid
HC: healthy control
kME: module eigenmetabolite connectivity, a measure of correlation between a metabolite and its assigned module
mPAP: mean pulmonary artery pressure
OCFA: odd-chain fatty acid
PAH: pulmonary arterial hypertension
PH: pulmonary hypertension
PVR: pulmonary vascular resistance
RHC: right heart catheterization
RV: right ventricle (or right ventricular)
RV-PV: right ventricular-pulmonary vascular
RVEF: right ventricular ejection fraction
TAPSE: tricuspid annular plane systolic excursion
WGCNA: weighted gene co-expression network analysis
WU: Wood units

## Supplemental Material

Supplemental Methods

Figure S1-S2

Tables S1-S5

Module Replication Atlas

PVDOMICS Module Results Atlas

## Novelty and Significance

What Is Known?

- Pulmonary arterial hypertension (PAH) is associated with widespread alterations in circulating metabolites, including abnormalities in lipid, amino acid, and steroid metabolism.
- Right ventricular (RV) function is the primary determinant of mortality in patients with PAH.
- Prior studies have linked individual circulating metabolites and pathways to RV function and outcomes in PAH.

What New Information Does This Article Contribute?

- The PAH metabolome is organized into reproducible, biologically coherent modules that represent structured intensification and reorganization of conserved metabolic programs rather than emergence of disease-specific pathways.
- Fatty acid oxidation and conjugated disposal modules are associated with adverse hemodynamics and worse RV function, whereas acylcholine and 5α-reduced androgen metabolite modules are associated with favorable right ventricular-pulmonary vascular phenotypes.
- Multicompartment gradient analysis reveals the pulmonary circulation as an active metabolic interface for steroid conversion, with transpulmonary uptake of sex hormone precursors and concurrent release of reduced androgen derivatives.

Metabolic dysregulation is a hallmark of pulmonary arterial hypertension (PAH), yet the organization of the circulating metabolome and its relationship to right ventricular–pulmonary vascular (RV-PV) function remain incompletely understood. This study applied weighted gene co-expression network analysis (WGCNA) to untargeted metabolomic data from 412 PAH patients in the multicenter PVDOMICS cohort to define this architecture. WGCNA identified 16 co-regulated metabolic modules organized around biologically coherent programs. A tripartite fatty acid axis spanning substrate pools, β-oxidation intermediates, and conjugated disposal products formed a central organizing structure, with downstream oxidation modules strongly associated with adverse hemodynamics and worse RV function. In contrast, acylcholine-enriched and 5α-reduced androgen metabolite modules were associated with favorable hemodynamic indices. Notably, module architecture was largely preserved in healthy controls, indicating that PAH involves reorganization of normal metabolic programs rather than emergence of novel pathways. Multicompartment sampling across pulmonary circulation revealed striking transpulmonary gradients for steroid modules, supporting the pulmonary vasculature as an active site of steroid conversion. Core modules and hub metabolites were recovered in an independent replication cohort. These findings provide a systems-level framework for understanding metabolic remodeling in PAH and identify coordinated metabolic programs that may serve as mechanistic targets and biomarkers of RV-PV dysfunction.

